# The stability flexibility tradeoff and the dark side of detail

**DOI:** 10.1101/2020.01.03.894014

**Authors:** Matthew R. Nassar, Vanessa Troiani

## Abstract

Learning in dynamic environments requires integrating over stable fluctuations to minimize the impact of noise (stability) but rapidly responding in the face of fundamental changes (flexibility). Achieving one of these goals often requires sacrificing the other to some degree, producing a stability-flexibility tradeoff. Individuals navigate this tradeoff in different ways, with some people learning rapidly (emphasizing flexibility) and others relying more heavily on historical information (emphasizing stability). Despite the prominence of such individual differences in learning tasks, the degree to which they relate to broader characteristics of real-world behavior or pathologies has not been well explored. Here we relate individual differences in learning behavior to self-report measures thought to collectively capture characteristics of the Autism spectrum. We show that that young adults who learn most slowly tend to integrate more effective samples into their beliefs about the world making them more robust to noise (more stability), but are more likely to integrate information from previous contexts (less flexibility). We show that individuals who report paying more *attention to detail* tend to use high flexibility and low stability information processing strategies. We demonstrate the robustness of this inverse relationship between *attention to detail* and formation of stable beliefs in a heterogeneous population of children that includes a high proportion of Autism diagnoses. Together, our results highlight that *attention to detail* reflects an information processing policy that comes with a substantial downside, namely the ability to integrate data to overcome environmental noise.

## Introduction

Successful decision making requires inferring important quantities such as the values and probabilities associated with potential decision outcomes through sequential observations over time. This inference process is difficult in changing environments, where optimal inference requires tracking the environmental statistics necessary to determine the most appropriate rate of learning (Behrens, Woolrich, Walton, & Rushworth, 2007; Browning, Behrens, Jocham, O’Reilly, & Bishop, 2015; McGuire, Nassar, Gold, & Kable, 2014; Nassar, McGuire, Ritz, & Kable, 2019b; Nassar et al., 2012; Prescott Adams & MacKay, 2007; Wilson, Nassar, & Gold, 2010; Yu & Dayan, 2005). In general, learning should be slow during periods of environmental stability in order to average over as many relevant observations as possible, but fast during periods of environmental change that render prior observations irrelevant to the problem of predicting future ones (Behrens et al., 2007; Browning et al., 2015; Nassar et al., 2016; Nassar, Wilson, Heasly, & Gold, 2010; Vaghi et al., 2017; Wilson, Nassar, & Gold, 2013). Human behavior, fMRI BOLD responses, and measures of physiological arousal display qualitative hallmarks of this sort of learning rate adjustment, suggesting that the brain implements meta-control over its own rate of learning in order to optimize behaviorally relevant inferences (Behrens et al., 2007; Browning et al., 2015; McGuire et al., 2014; Nassar et al., 2012; Nassar, McGuire, Ritz, & Kable, 2019b; Prescott Adams & MacKay, 2007; Wilson et al., 2010; Yu & Dayan, 2005).

However, learning rate, and adjustments thereof, differ dramatically across individuals, age groups, and clinical populations (Behrens et al., 2007; Browning et al., 2015; Nassar et al., 2010; 2016; Vaghi et al., 2017; Wilson et al., 2013). Some individuals tend to adjust beliefs rapidly irrespective of environmental statistics, leading to flexible but unstable beliefs, whereas others tend to adjust more slowly giving rise to inflexible but stable beliefs (Nassar et al., 2010). In principle, such differences might arise through learning about environmental statistics over a much longer time course, such as over development (Nassar et al., 2016) or even evolution (Krugel, Biele, Mohr, Li, & Heekeren, 2009; Stein, Newman, Savitz, & Ramesar, 2006). This longer timescale meta-learning might in some cases appropriately bias an individual towards one end of the stability/flexibility spectrum; however in other cases it could potentially go awry and give rise to pathological belief updating. For example, recent work has suggested individuals with obsessive compulsive disorder tend to over-learn from new information (Vaghi et al., 2017), limiting stability of beliefs. Consistent with a prominent theory of autism (Sinha et al., 2014), similar conclusions have been made about autistic individuals under some conditions (Lawson, Mathys, & Rees, 2017), although other studies have failed to identify differences between autistic individuals and controls (Manning, Kilner, Neil, Karaminis, & Pellicano, 2016).

These mixed results may result in part from heterogeneity within the autism spectrum. Autism is a broad diagnostic category characterized by deficits in social communication as well as restricted and repetitive patterns of behavior (RRBs). RRBs include inflexible adherence to routines, inflexibility to changing contexts, rigid thinking patterns, and increased *attention to detail*. Although the neural origins of an increased focus on details remains unknown, it has been described colloquially as “missing the forest for the trees” and theoretically as “weak central coherence” (Frith, 1989; Happé & Frith, 2006), “enhanced discrimination and reduced generalization” (Plaisted, 2001), and “enhanced perceptual functioning” (Mottron, Dawson, Soulières, Hubert, & Burack, 2006). Studies investigating increased *attention to detail* tend to employ experimental tasks that require an individual to extract smaller feature from a larger whole (i.e. Navon figures, block design task, embedded figures), in which performance can indicate a preference or bias towards local features or the more global whole. Several studies have found individuals with autism spectrum disorder (ASD) to have a local perceptual bias (Dakin & Frith, 2005; Happé, 1999; Simmons et al., 2009). Such a local bias, if extended to learning through time, might predict an over-reliance on recent information (supporting flexibility), at the expense of integration over relevant historical information (limiting stability).

One factor limiting much of the previous research on *attention to detail* is the focus on dichotomous groups of individuals with or without an ASD diagnosis, a study design that does not take into account that behavioral manifestations of ASD are heterogeneous within the disorder and also are present in the general population. Importantly, local/global perception has also been shown to vary in the general population (Dale & Arnell, 2013; McKone et al., 2010; Scherf, Behrmann, Kimchi, & Luna, 2009). Recent work has shown that quantitative traits of autism measured both in the general population and within clinically-diagnosed cohorts are associated with the ability to disembed a smaller figure from a larger shape (Sabatino DiCriscio & Troiani, 2017; 2018). Further, it has been shown that the ability to disembed a local part from a global whole are not present in every individual with ASD, indicating that measuring trait dimensions is important in heterogeneous disorders like autism (DiCriscio, Hu, & Troiani, 2019).

Here we use a trait dimension approach to examine the relationship between *attention to detail*, a prominent feature of autism, and the degree to which individuals implement learning policies favoring either stability or flexibility. We relate individual differences in learning behavior (stability/flexibility tradeoff) to a quantitative measure of autism traits (Autism Spectrum Quotient; AQ), designed to capture characteristics of autism that extend outside of traditional diagnostic boundaries. We examine this in two separate populations: healthy young adults and children with a range of developmental abilities, including autism. We show that the young adults who update beliefs the least in the face of conflicting information integrate more effective samples into their beliefs about the world, making them more robust to noise (more stability), but also are more likely to integrate information from previous contexts (less flexibility). The individuals who show the opposite pattern of results (high flexibility/low stability), tended to score higher on the *attention to detail* subscale of the AQ. We confirm this inverse relationship between *attention to detail* and formation of stable beliefs in a population of children that includes a high proportion of clinical autism diagnoses. Together, our results highlight that *attention to detail* reflects an information processing policy that comes with a substantial downside, namely the inability to integrate data to overcome environmental noise.

## Methods

### Subject populations

#### Experiment 1

43 young adults (20 female, mean[std] age = 2.4[3.4] years, Mean[std] WASI FSIQ = 112[10.4]) were recruited from a local community population to participate in our first behavioral study.

#### Experiment 2

37 children (17 female, mean[std] age = 9.5 [2.5]) were recruited to participate in our second behavioral study. In order to obtain a range of autism traits in the sample, we identified participants using a broad recruitment strategy. This included identifying participants based on patient referral to a neurodevelopmental clinic in Lewisburg, Pennsylvania, as well as from health system wide advertisement and the surrounding community. On the day of research testing, all participants completed a cognitive assessment to document IQ (WASI-II: Wechsler abbreviated scale of intelligence, 2nd edition; Wechsler, 2011). If an IQ test was ascertained as part of their clinic appointment that day, we used the clinically ascertained IQ score. All participants assented to protocols approved by the institutional review board (IRB) at the authors’ home institution. Twelve of our participants had a clinical diagnosis of autism or ASD based on assessment by our neurodevelopmental pediatricians and support staff.

### Study Session

Each experimental session involved performing a computerized predictive inference task (Nassar et al., 2016; Nassar, Bruckner, & Frank, 2019a), completing the Autism Quotient questionnaire, and a cognitive assessment (WASI FSIQ).

### Predictive inference task

Each participant completed a computerized predictive inference task that required them to infer the location of an unobservable helicopter based on the locations of bags that had previously fallen from it (McGuire et al., 2014). The task included two conditions that favor different adaptive learning strategies (d’Acremont & Bossaerts, 2016; Nassar, Bruckner, & Frank, 2019a). In one condition the helicopter was generally stationary but occasionally underwent “changepoints” at which its position was reset to a random horizontal position on the visible screen, in the other condition the helicopter “drifted” slightly from trial-to-trial. On each trial a bag would fall from the top of the screen, providing the participant with some information about the helicopter location. In the changepoint condition this information was always relevant – as bag locations were normally distributed around the helicopter position. However, in the condition with the drifting helicopter, bags were occasionally sampled from a uniform distribution extending across the entire screen, giving rise to “oddball” events that were unrelated to the true helicopter location. Both conditions were fully instructed using an extended training period in which the true helicopter position was visible to the participants. On each task trial, participants were asked to move a bucket to the inferred location of the helicopter in order to catch tokens in the bags that were translated into points.

### Autism Spectrum Quotient (AQ)

The AQ is a self-report measure aimed to assess ASD-like traits across the general population (Baron-Cohen, Wheelwright, Skinner, Martin, & Clubley, 2001). This measure assesses five trait domains, including communication, social skills, attention switching, imagination, and *attention to detail*. Using a 4-point Likert scale, a participant responds with how strongly they agree or disagree with a given statement. Each item is scored based on whether a given trait is endorsed, with half of the items requiring an *agree* and half requiring a *disagree* response to endorse an ASD-like trait. Item scores are summed to generate both a total score as well as subscale scores. In Experiment 1, young adults completed a self-report version of the AQ, while in Experiment 2, parents completed a parental-report version of the AQ on their child’s behavior.

### Subject Exclusion

For both studies, subjects were excluded if they did not meet a basic performance standard designed to determine whether they were actually attempting to complete the task (mean distance between bucket and helicopter position of less than 45 units). This performance standard was met by all participants in the young adult population but did lead to exclusion of one participant in the developmental cohort (see figure S2). In addition, 8 participants in the developmental cohort did not complete the AQ due to time constraints, and thus were not included in the correlations between AQ measures and task performance. After participant exclusion, our developmental cohort included 29 participants, 11 of whom had an autism diagnosis.

### Normative learning model

Normative learning was assessed using a reduced Bayesian model that has been described previously for the changepoint (Nassar et al., 2010) and oddball (Nassar, Bruckner, & Frank, 2019a) conditions.

### Single trial learning rates

Participant bucket positions and computer generated bag locations were used to compute trial-by-trial prediction errors (the difference between bag location and the center of the bucket on a given trial) and prediction updates (the bucket location on a subsequent trial minus the bucket location on the current trial). In order to estimate the degree of influence of each bag on the subsequent behavior of the participant, we computed a single trial learning rate by dividing the update made on each trial by the prediction error observed on that trial (Nassar et al., 2010). Learning rates computed in this way that were greater than 1 or less than 0 were set to 1 or 0, respectively. Single trial learning rates were categorized into three groups: 1) total updates [>.8], 2) moderate updates [.2 to .8], and 3) non-updates (<.2).

### Characterizing the content of participant beliefs

To better understand how the exact sequence of learning rates employed by each participants affected the precision and flexibility of their beliefs, we re-represented participant beliefs (bucket position) on each trial as a weighted mixture of previous outcomes (bag locations). To do so, we stepped through the sequence of single trial learning rates and for each trial 1) assigned weight to the newest outcome in proportion to the learning rate on that trial, and 2) updated the weight assigned to all previous outcomes by multiplying their weight (computed on the previous trial) by one minus the current trials learning rate. This procedure produced a vector of weights that, when multiplied by the corresponding vector of outcomes, resulted in the exact belief of the participant.

In order to assess the flexibility of beliefs, we quantified the proportion of the weight profile that was attributed to relevant outcomes. In the changepoint condition, relevant outcomes were defined as those having occurred since the most recent changepoint (eg. bags that fell from the current helicopter location). In the oddball condition, all non-oddball outcomes were considered to be relevant (eg. bags normally distributed around helicopter).

In order to assess precision beliefs, we quantified the effective number of outcomes from which they were composed. Specifically, we computed effective samples as follows:

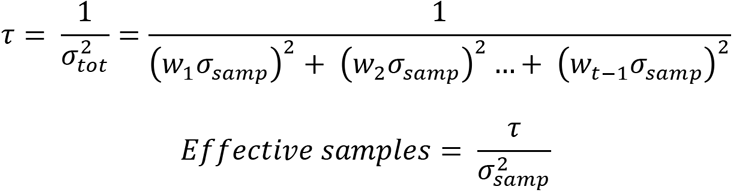

where *τ* reflects the precision (inverse variance) of beliefs, 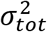 is the variance on the weighted mean of samples, 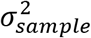 is the variance on each sample, and *w* reflects the weight given to that sample during updating. Our effective samples measure simply normalized the belief precision in terms of the precision of a belief based on a single observation.

### Statistical analysis

Rank order correlations between task measures and AQ sub-scale measures were computed using Spearman’s Rho. Linear regression was used for followup analyses designed to statistically control for other factors including IQ, age, and gender. All analyses and models were implemented in Matlab (The MathWorks, Natick, MA) and all code and anonymized data will be made available upon publication on the corresponding authors website (https://sites.brown.edu/mattlab/resources/).

## Results

### Experiment 1

Young adults made predictive inferences in both changepoint and oddball contexts. Participants specified predictions about the location of an unobservable helicopter (Fig 1A, prediction panel) in order to catch bags (Fig 1A, outcome panel). Predictions were updated on each trial (Fig 1A, update panel) according to the most recently observed bag location, and knowledge of the underlying generative structure (changepoint/oddball). In the changepoint condition, normative learning (Fig 1B, pink line) prescribed rapid updating in response to unexpected bag locations, as these outcomes were likely associated with a change in the helicopter location. In contrast, in the oddball condition normative learning (Fig1B, pink) required ignoring unexpected bag locations, which were likely to be oddballs unrelated to the actual helicopter position. Predictions made by an example participant (Fig1B&C, blue) conform well to normative model predictions. The normative learning model adjusts learning rate from trial to trial according to the probability that the observed outcome reflects a changepoint (Fig 1D, orange) or oddball (Fig 1E, orange), depending on the current task condition, as well as an estimate of uncertainty about the current helicopter location (Fig 1D&E, yellow; (Nassar et al., 2012)).

**Figure 1:**
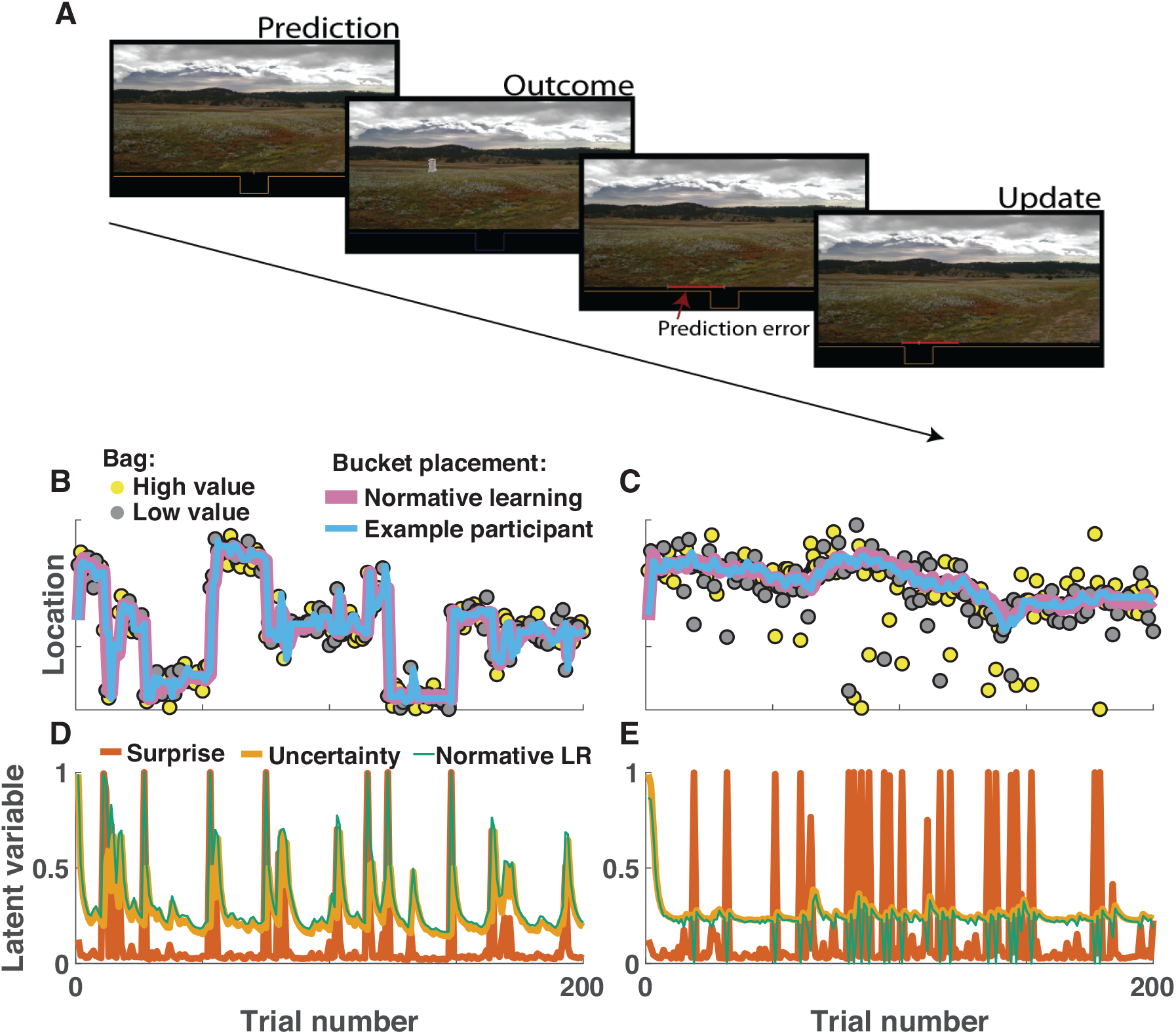
Predictive inference task measures learning in different statistical contexts. **A)** On each trial, participants were required to adjust the position of a bucket to catch bags of coins that would be dropped from an unobservable helicopter. Subjects were not able to observe the helicopter, and thus forced to use the history of bag locations and knowledge about the environmental statistics to inform bucket placement. **B-C)** Example data from a single subject performing the predictive inference task in changepoint (**B**) and oddball (**C**) conditions. **B**) In the changepoint condition, the helicopter (not shown) remained in a single screen position (ordinate) for a number of trials (abscissa), before occasionally relocating to a new screen position (changepoint). Bag locations (yellow and gray points) were drawn from a normal distribution centered on the helicopter location. Inferences about the helicopter location made by a normative learning model (pink line) and bucket placements made by an example subject (blue line) are both rapid to adjust after changepoints in the helicopter location. **C**) In the oddball condition, the helicopter position drifted slowly from one trial to the next, and bag positions were either drawn from a normal distribution centered on the helicopter location (90% of trials) or a uniform distribution across the entire task space (10% of trials). **D&E**) The normative learning model adjusted learning rate (green line) on each trial according to uncertainty (yellow) and surprise (orange). In the changepoint condition **(D)** surprise was indicative of changepoints and increased learning rates, whereas in the oddball condition **(E)** surprise was indicative of an uninformative oddball and thus promoted lower learning rates.

The distribution of single trial learning rates used by participants differed across the two conditions, qualitatively in accordance with the normative predictions. On changepoint trials, participants tended to use high learning rates (Fig 2A), whereas on oddball trials where bag locations were equally surprising but unrelated to the true helicopter location, participants tended to use learning rates near zero (Fig 2B). Distribution of learning rates across unsurprising trials tended to be more similar across the conditions, with a fair number of small and moderate learning rates employed (Fig 2C&D). These relative patterns of learning were consistent across subjects, with total-updates (learning rate > 0.8) decreasing with increasing trials after a changepoint (Fig 2E; linear effect of trials after changepoint on total updating: t = −8.0, df = 42, p=5×10^−10^) and non-updates (learning rate < 0.2) elevated on oddball trials (Fig 2F; contrast non-updating on oddball versus other trials: t = 2.9, df = 42, p=0.007).

**Figure 2:**
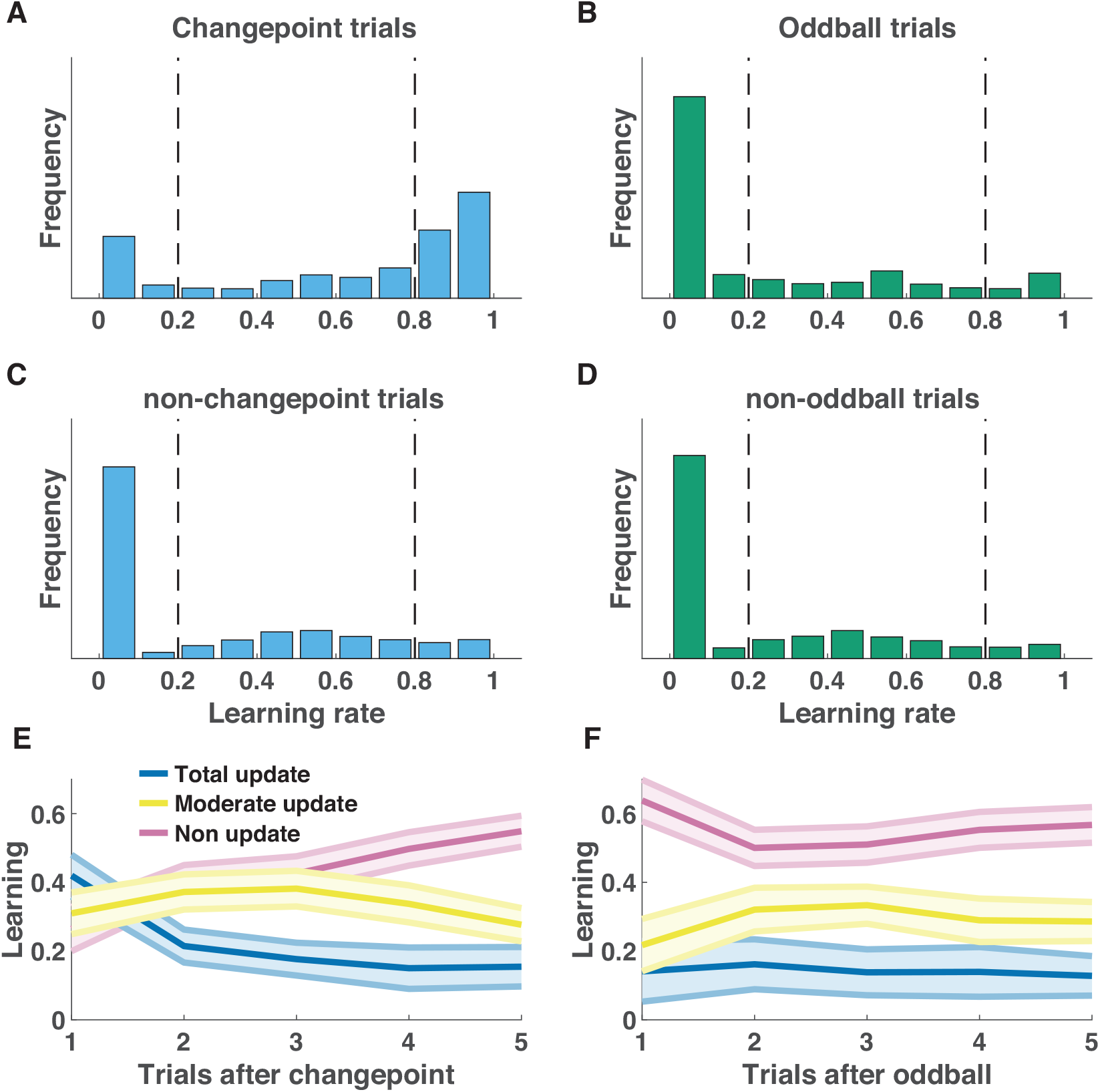
Participant learning rates were sensitive to task condition and surprising outcomes. **A-D)** Single trial learning rate frequency histograms for changepoint (**A**) and oddball (**B**) trials, as well as for non-changepoint (**C**) and non-oddball (**D**) trials. Single trial learning rates are categorized into three types: non updates, moderate updates, and total updates, depending on their value with respect to criterion values (dotted vertical lines). **E-F**) Mean/SEM proportion of each category of learning rates used as a function of time since the previous surprising event [changepoint (**E**) or oddball (**F**)].

Despite the preservation of relative learning patterns, participants differed markedly in their overall learning rate distributions, with some participants almost never making a total update and others using total updates on approximately six out of ten trials. In principle, these differences could reflect different policies toward optimizing either the stability or flexibility of beliefs. In order to test whether such a stability/flexibility tradeoff exists, we examined how individual differences in total update frequency related to performance in each of the task conditions. In the changepoint condition, higher total update frequency tended to be associated with smaller errors on changepoint trials (Figure 3A; Spearman’s rho = −0.34, p = 0.02) but larger errors during periods of stability (Figure 3C; Spearman’s rho = 0.66, p = 1.5 × 10^−6^), supporting the idea that individuals may differ in their relative concern for stability versus flexibility of beliefs. Performance in the oddball condition, in contrast, tended to favor more stable belief updating strategies, with more frequent total updates leading to worse performance on oddball trials (Fig 3B&D; Spearman’s rho = 0.66, p = 1.5 × 10^−6^). Thus, individual differences in learning, specifically the frequency of total updating, predicted individual differences in performance in a manner that suggests different policies regarding toward optimizing stability or flexibility.

**Figure 3:**
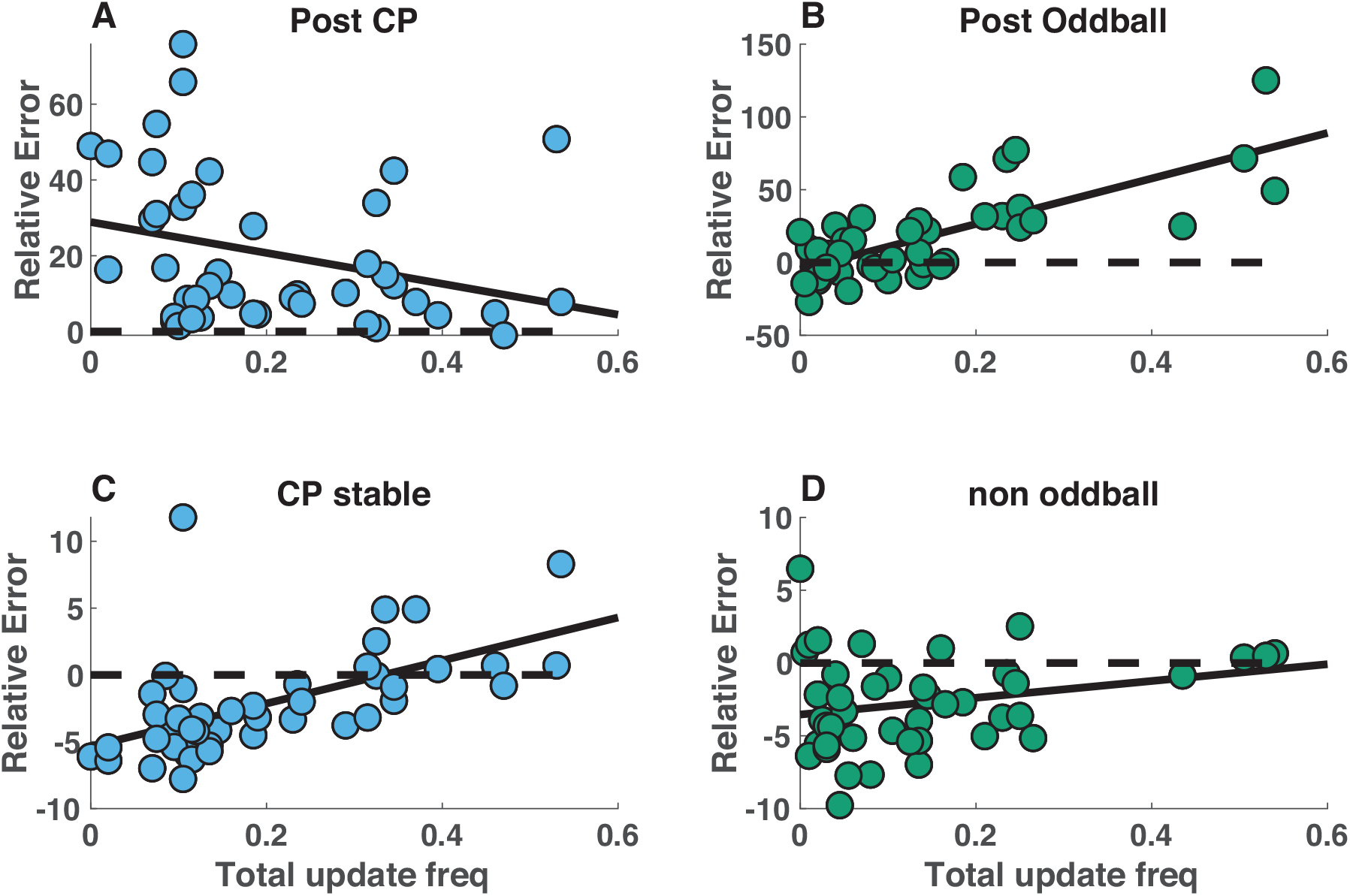
Individual differences in performance were driven by individual differences in the frequency of total updates. **A-D)** Relative error magnitude (ordinate) is plotted against frequency of total updating (abscissa) for each individual subject (points) in the two task conditions (changepoint = blue, oddball = green). Top panels reflect performance on the trial subsequent to a surprising outcome (**A** = post-changepoint, **B** = post-oddball) whereas bottom panels reflect performance during periods of stability (>5 trials after most recent changepoint (**C**) or oddball (**D**)). Relative error would be zero if participants were using only the most recent relevant outcome in order to make their prediction (dotted line) and thus achieving negative relative errors requires using bucket placements that integrate information from more than one previous outcome.

In order to more explicitly test for individual differences in the stability and flexibility of beliefs, we used the sequence of learning rates preceding each prediction to determine the weighted contribution of each previous outcome to that prediction (see methods). When applied to simple fixed learning rate models, this method revealed the expected exponential decay of weight across previous outcomes, with higher learning rates corresponding to higher rates of decay (Fig4A, blue&yellow). Normative learning relies on weights with more complex dynamics, which are approximately uniform across trials since the most recent changepoint, but zero on trials prior to the most recent changepoint (Fig4A, green). The lack of weight attributed to outcomes prior to the previous changepoint affords the normative model flexibility, whereas the roughly uniform weighting of outcomes since the previous changepoint facilitates precise beliefs by averaging over the noise associated with each relevant outcome. This benefit in precision can be quantified through the number of effective samples comprising the current prediction, which grows nearly linearly for the normative model during periods of stability but rapidly decays to one after a changepoint (Fig 4b, yellow). Note that a simple high fixed learning rate model, which is flexible in that it rapidly discards old and potentially irrelevant information (Fig 4a, blue), never accumulates even two effective samples (Fig 4b, blue), and is therefore highly sensitive to noise. This illustrates the stability flexibility tradeoff – rapid learning can promote flexibility at the expense of precision during periods of stability.

**Figure 4:**
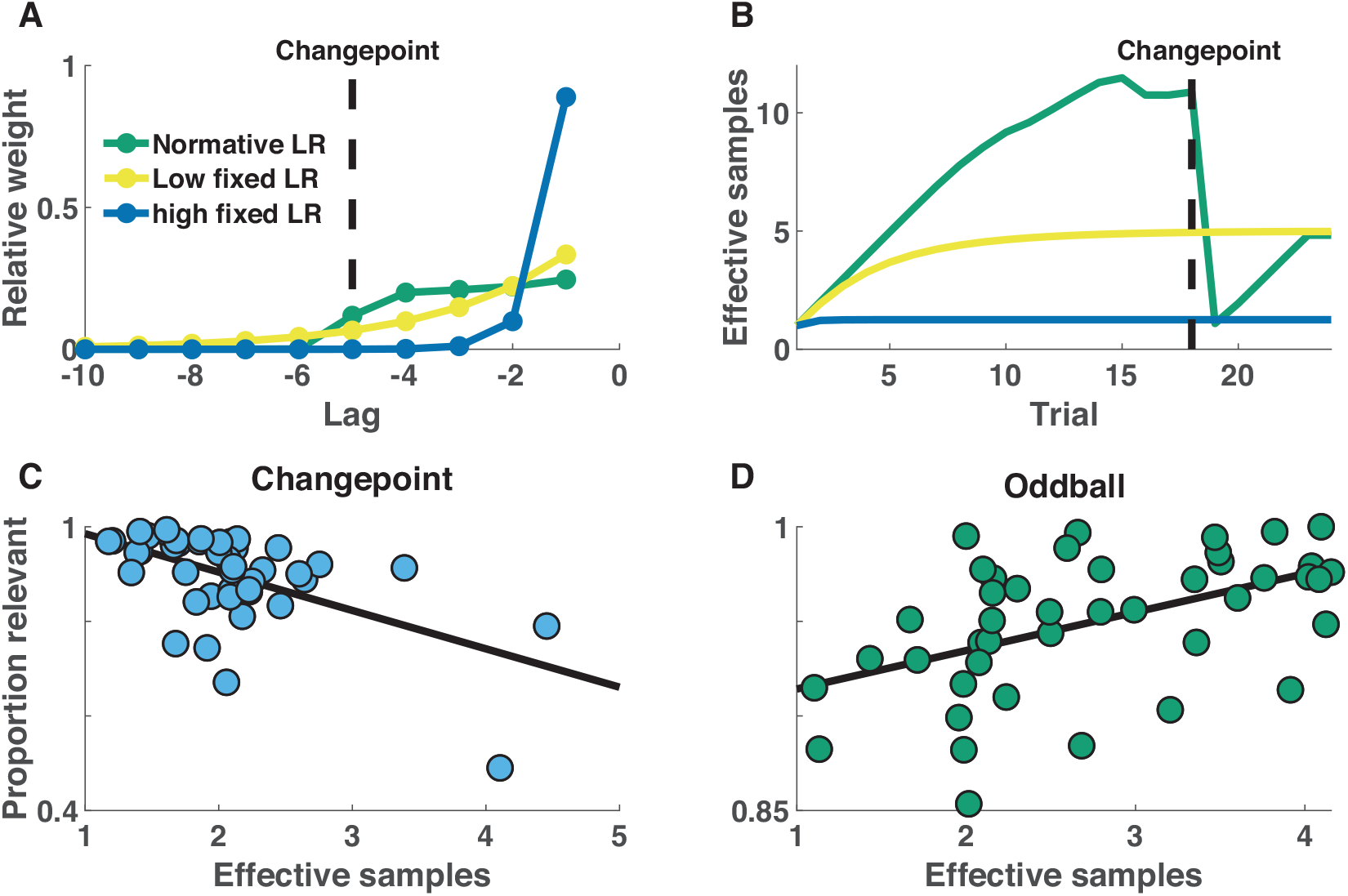
Individual differences in performance are attributable to a fundamental tradeoff in the quantity and relevance of samples from which a belief is composed. **A)** Bucket placements on a given task trial can be decomposed into a weighted average of previous observations. Higher learning rates correspond to a greater proportion of weight attributed to recent observations (compare blue and yellow lines) and normative learning approximates a flat weighting of all observations since the previous changepoint (green). Observations occurring prior to the most recent changepoint are irrelevant to the inference process, and thus the proportion of weights attributed to observations occurring since the last changepoint quantifies the relevance of samples from which the belief is composed. **B)** For a bucket placement on a given trial, the distribution of weights over previous observations can be used to infer the effective number of samples incorporated into that belief (which scales with the precision – or inverse variance – of that belief). High learning rate models, which rely predominantly on the most recent observations, rely on beliefs with the fewest effective samples (blue). Normative learning approximates linear growth of effective samples during periods of stability, but rapid collapse of effective samples after observing a changepoint (green). **C-D)** Participants who incorporated the most samples into their beliefs (abscissa) tended to rely on less relevant information (ordinate) in the change-point condition (**C**), whereas this relationship reversed in the oddball condition (**D**).

The stability flexibility tradeoff was also evident from individual differences in our task. Individuals who were most flexible (eg. had a high proportion of weight associated to relevant outcomes) tended to base predictions on fewer samples (Fig 4C; Spearman’s rho = −0.49, p = 9.1 × 10^−4^) in the changepoint condition. Conversely, individuals who incorporated more outcomes into their predictions, tended to include a higher proportion of irrelevant outcomes, making their predictions less flexible in the face of changepoints. In the oddball condition, where task-relevance was unrelated to recency, this relationship reversed such that participants who incorporated the most samples, also tended to have the highest proportion of relevant ones (Fig4d; Spearman’s rho = 0.53, p = 3.1 × 10^−4^). Taken together, these results suggest that individuals differ in their relative emphasis on the precision of beliefs, or their flexibility in the face of changing contexts.

An important motivating question of this work was to examine the degree to which such differences in stability/flexibility policy might relate to broader patterns of real-world behavior, with respect to traits that are elevated in ASD. In line with this idea, healthy young adults who scored highest on the *attention to detail* sub-domain of the AQ incorporated fewer effective samples into beliefs in the changepoint condition (Fig 5A; Spearman’s rho = −0.43, p = 0.005), but those samples were more relevant (Fig 5C; Spearman’s rho = 0.33, p = 0.03). No relationships between these measures of stability and flexibility were observed in relation to other sub-domains of the AQ (Fig S1; all 8 p values > 0.1) or in relation to IQ (p = 0.16, 0.28). There was a trend toward the same negative relationship between precision and *attention to detail* in the oddball condition (Spearman’s rho = −0.30, p = 0.06), however the advantage of high *attention to detail* individuals in terms of sample relevance was not apparent in this condition (Spearman’s rho = −0.22, p = 0.17). Taken together, these results suggest “*attention to detail*”, one specific aspect of behavioral variability that has been associated with ASD, directly relates to stability/flexibility policy, with individuals higher on “*attention to detail*” favoring flexibility even at the expense of stability.

**Figure 5:**
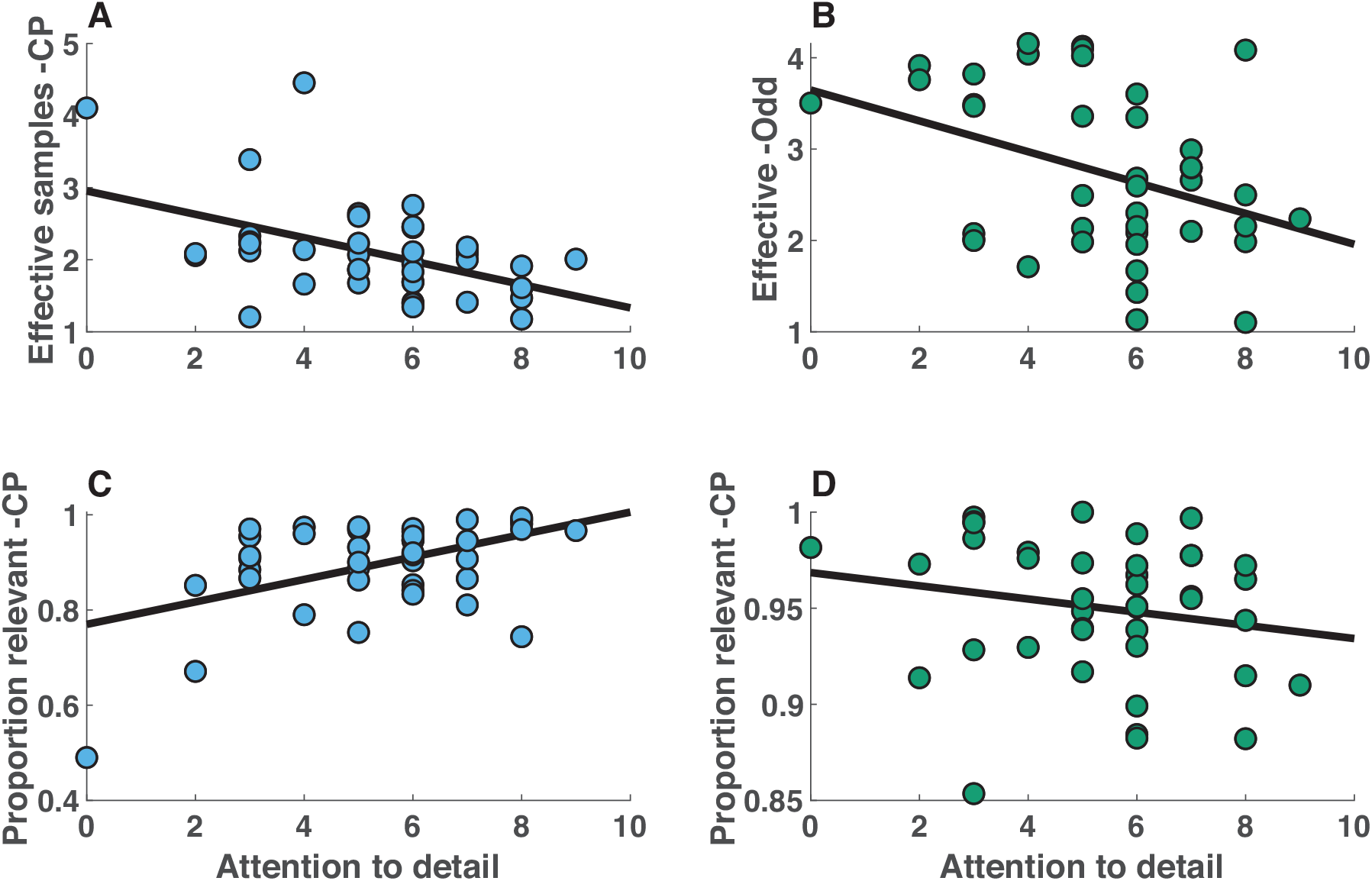
*Attention to detail* predicts individual differences in stability/flexibility policy. **A-B)** Individual differences in the effective sample size of beliefs (ordinate) were negatively related to self reported scores on the *attention to detail* subscale of the Autism Spectrum Questionnaire (abscissa) in both changepoint (**A**) and oddball (**B**) conditions. **C-D)** Individual differences in the proportion of samples attributed to relevant observations (ordinate) were positively related to self reported scores on the *attention to detail* subscale of the Autism Spectrum Questionnaire (abscissa) in the changepoint (**A**) but not the oddball (**B**) conditions.

### Experiment 2

In order to test the generality of the relationship between *attention to detail* and stability/flexibility and to examine it across a wider range of behavioral phenotypes that includes individuals with ASD, we conducted a second behavioral study in a heterogenous population of children (N = 37, mean[STD] age = 9[2.5], 17 female). The group included 12 participants diagnosed with ASD, as well as 25 children recruited from the local community.

In general, behavior of the children included far fewer updates than that of the young adult population. Non-updates were the most common updating category, even on changepoint trials that should require total updates (Figure 6). In principle, non-updates could limit flexibility by reducing responsiveness to new information after a changepoint, but could also limit precision of beliefs during periods of stability by preventing incorporation of new information into existing beliefs. Consistent with this idea, there was no evidence for a stability flexibility tradeoff in either condition for the developmental cohort (Figure 7A; Spearman’s Rho 0.21, p = 0.2 for the changepoint condition and Rho −0.03, p = 0.87 for the oddball condition).

**Figure 6:**
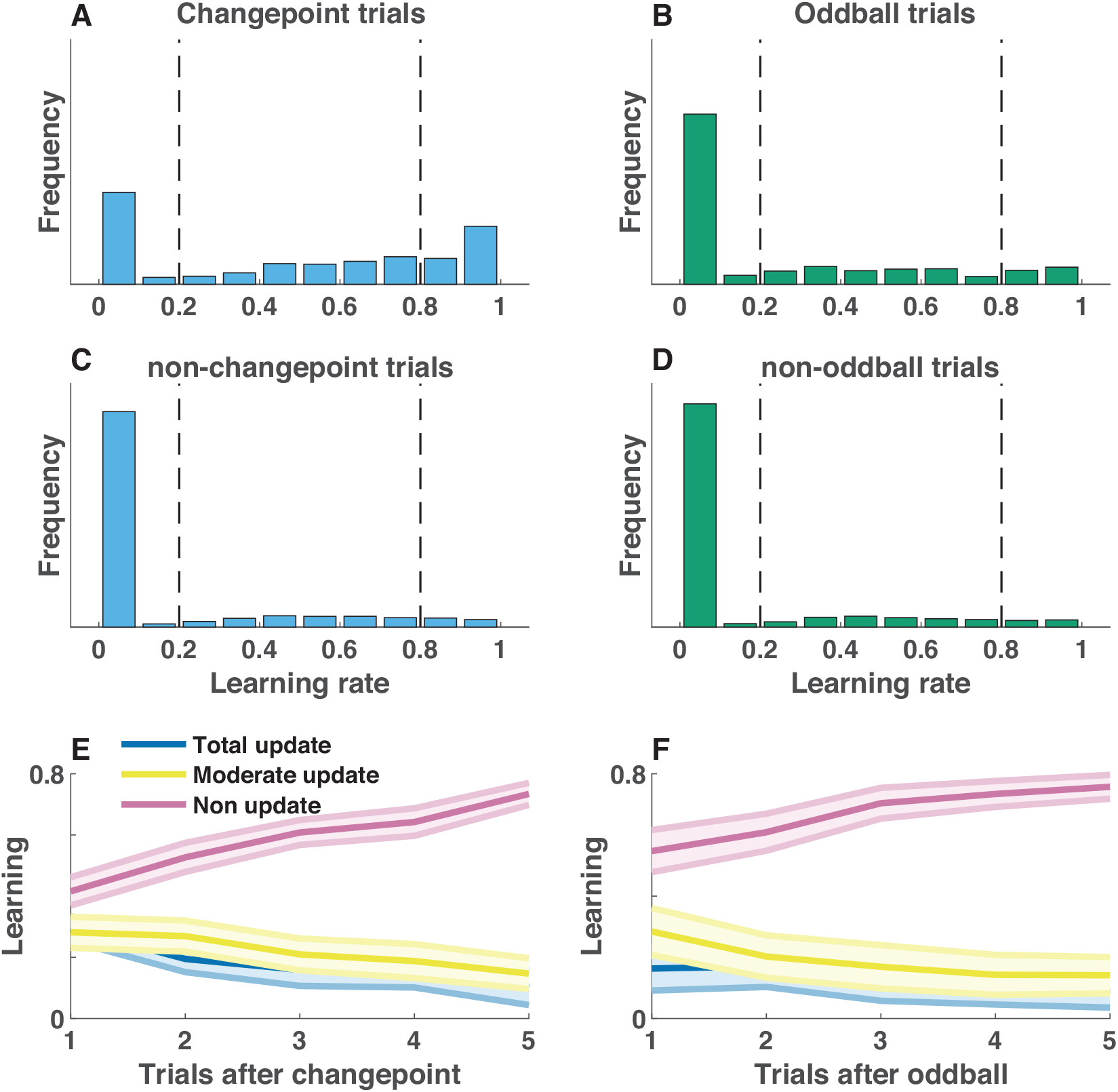
Non-updates are increased and condition-differences are less pronounced in a heterogeneous population of children. **A-D)** Single trial learning rate frequency histograms for changepoint (**A**) and oddball (**B**) trials, as well as for non-changepoint (**C**) and non-oddball (**D**) trials. Single trial learning rates are categorized into three types: non updates, moderate updates, and total updates, depending on their value with respect to criterion values (dotted vertical lines). **E-F**) Mean/SEM proportion of each category of learning rates used as a function of time since the previous surprising event [changepoint (**E**) or oddball (**F**)].

**Figure 7:**
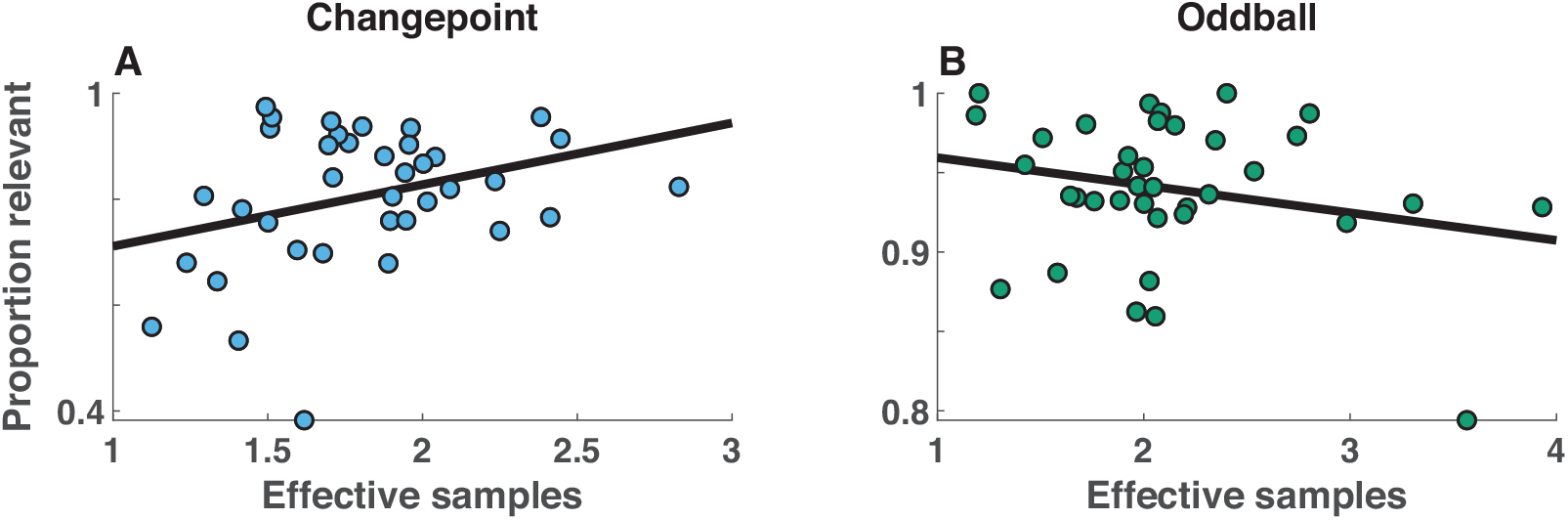
Stability/flexibility tradeoff does not explain individual differences in updating among a heterogeneous population of children. **A-B)** For each participant (points), the mean proportion of samples composing the belief that are relevant to the current statistical context (abscissa) is plotted against the total number of effective samples composing the belief (ordinate) separately for changepoint (**A**) and oddball (**B**) conditions. Trend lines indicate least squares fit to data.

Despite the lack of evidence for a stability flexibility tradeoff in this heterogenous population of children, *attention to detail* was still related across participants to lower precision beliefs. The average number of effective samples in participant beliefs aggregated across conditions was greatest for individuals with the lowest *attention to detail* scores (Spearman’s Rho = −0.50, p = 0.006). This relationship was similar in the two conditions (Fig 8A&B), but only reached statistical significance in the oddball condition (Spearman’s Rho for CP and ODD conditions: −0.25, −0.56; p values: 0.19, 0.001). Unlike in the young adult population, *attention to detail* did not confer any advantage in terms of flexibility to children (Fig 8C&D; p-value for correlations in both conditions > 0.5), likely due to additional variance in flexibility measures attributable to non-updating at changepoints (Fig 6E, pink). The relationship between *attention to detail* and belief precision was robust to inclusion of IQ, age, and gender into the explanatory model (Mean[95% conf int] beta for extended regression model = −0.20[−0.33, −0.06], t = −3.0, dof = 24, p = 0.006). Taken together, these results suggest that *attention to detail* comes at a significant cost to the precision of beliefs, whereas the potential benefits of *attention to detail* in terms of belief flexibility are population dependent.

**Figure 8:**
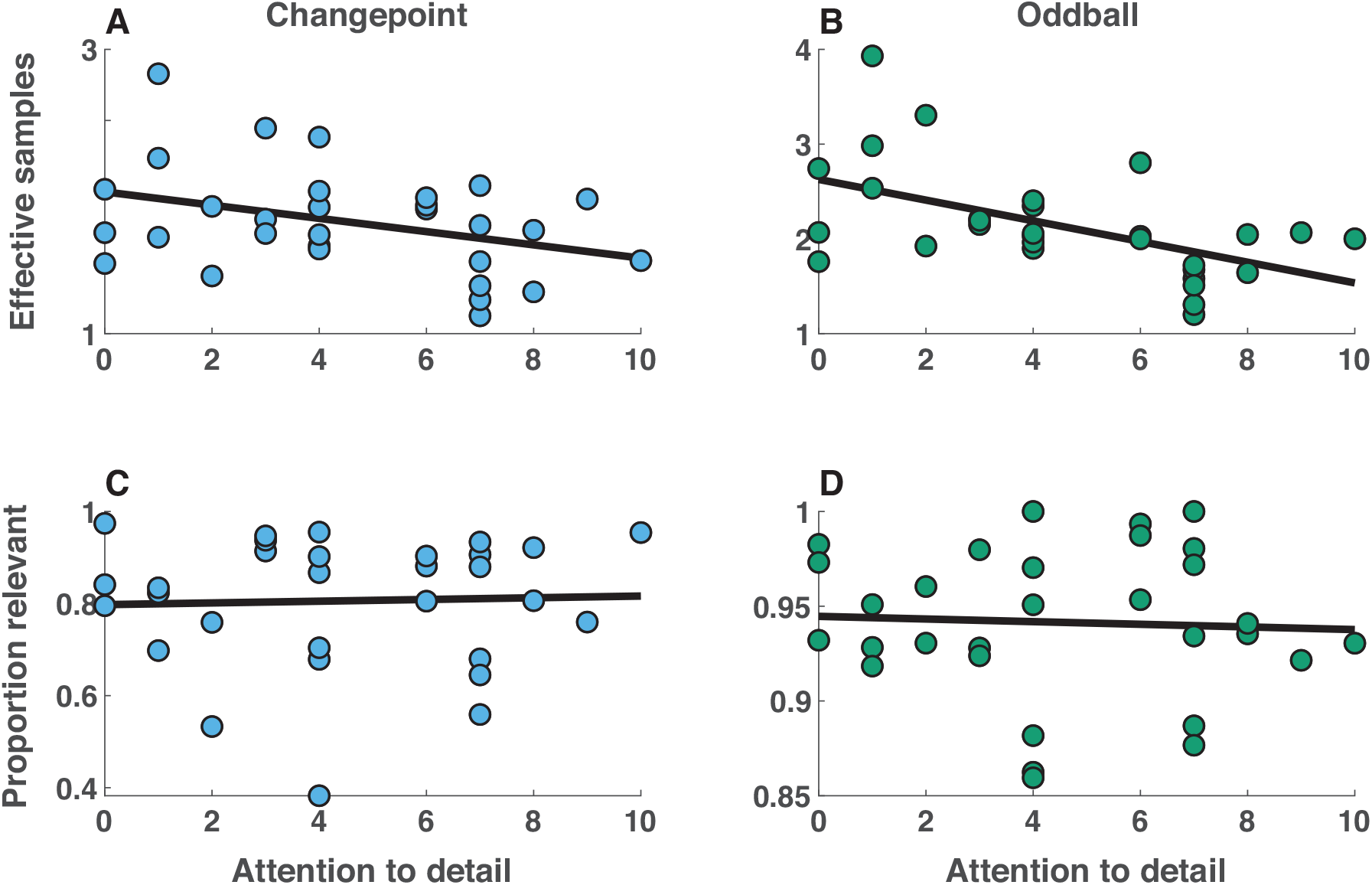
*Attention to detail* is inversely related to the number of effective samples composing beliefs in a heterogeneous population of children. **A-B)** For each participant (points), the total number of effective samples composing the belief (abscissa) is plotted against self reported scores on the *attention to detail* subscale of the AQ (ordinate) revealing a negative relationship in both changepoint (**A**) and oddball (**B**) conditions. **C-D)** In contrast, the proportion of relevant samples (abscissa) was unrelated to *attention to detail* scores (ordinate) for both changepoint (**A**) and oddball (**B**) conditions.

## Discussion

Autism is a multi-dimensional construct with a broad behavioral profile. One autism-linked dimension, *attention to detail*, has been related to a focus on local, as opposed to global, stimulus information. Here we explored whether this local bias might exist in time as well as in space, and whether such a bias would manifest in highly flexible but unstable beliefs. We confirmed high *attention to detail* young adults were more prone to completely updating beliefs in the face of contradictory information, and that this led them to form beliefs that were more flexible but which incorporated fewer observations, and thereby less robust to noise during periods of stability. We replicated the negative relationship between *attention to detail* and belief precision in a developmental cohort that included both typically developing and children with autism, but did not identify any advantages of higher attentional to detail individuals in this population with respect to flexibility. Taken together, our results highlight that high *attention to detail*, a prominent feature of autism, has profound implications for the way that information is used over time – promoting the use of recent, rather than historical information, and limiting the degree to which beliefs integrate over multiple observations.

To a first approximation, our results are consistent with basic tenants of “Weak Central Coherence Theory” (Frith, 1989; Happé & Frith, 2006). Specifically, the “global” aspect of our task might be considered to be the entire sequence of bag locations falling from the current helicopter location, whereas the “local” aspect might be considered to be the most recent bag location. We found that individuals who are high on *attention to detail*, a trait sometimes associated with autism, tend to focus on temporally local information, and form beliefs that incorporate fewer samples from the “global” category. This work nicely parallels work in the perceptual domain that has defined local and global in terms of space (O’Riordan & Plaisted, 2001; Plaisted, Dobler, Bell, & Davis, 2006; Sabatino DiCriscio & Troiani, 2017; Suzanne Scherf, Luna, Kimchi, Minshew, & Behrmann, 2008). However, our work did not focus on the binary autism distinction, but rather directly linked to measures of *attention to detail*. This is only one of many traits that is prevalent in ASD and indeed, in our populations was only very minimally related to other autism-linked traits (See supplementary figures 1&3). Our focus on traits might have heightened our ability to see such an effect, where other recent work that has compared autism to controls has had mixed results (Lawson et al., 2017; Manning et al., 2016).

Our results also speak to the more general tradeoff that the brain faces with respect to controlling the use of recent versus historical information. In stable regimes, optimizing this tradeoff requires integrating over all relevant historical observations, but changes in the environment require rapidly refocusing on recent observations to afford flexibility. We found, as had been reported previously, dynamic adjustments in the use of information according to environmental statistics, but we also noted an extremely wide range of overall learning behaviors (Figure 3). Note that this need not be the case from a computational perspective; participants were trained explicitly on the generative structure of the task, and had more than enough experience to estimate the rate of changepoints and oddballs, were they inferring these meta-parameters from the task observations (Nassar, Bruckner, & Frank, 2019a; Wilson et al., 2010). Thus, if participants came into the task without strong predispositions towards favoring either stability or flexibility, then they should have all arrived at similar policies by the end of the training session. However, this is not what we observed. One might argue that the heterogeneity across the individual participants reflects completely different task strategies, however the link between *attention to detail* and learning policy observed in our two experiments (Figures 5&8) suggests that participants come to the task with a systematic predisposition toward a specific learning strategy, either favoring the use of recent information for flexibility (high *attention to detail*) or favoring the integration of data over time for stability (low *attention to detail*).

One interesting question stemming from this work relates to the developmental timescale and origin of these predispositions from a neural perspective. For example, many aspects of RRBs present in ASD are considered age appropriate for most neurotypical toddlers (i.e. insistence on sameness, circumscribed/hyperfocused interests, inflexibility). One reason these behaviors are considered atypical in ASD is that they persist in older children and adults with the diagnosis, significantly contributing to impairment in real-world situations. From the neural perspective, the heterogenous behavioral traits of ASD are not thought to stem from specific regions of the brain, but rather from atypical connectivity between brain regions. Atypical connectivity has been identified in numerous studies across multiple brain networks in ASD using various neuroimaging methods, including structural and functional MRI, EEG and MEG, and fNIRS (for reviews, see Hull et al., 2017; O’Reilly, Lewis, & Elsabbagh, 2017; Rane et al., 2015; Zhang & Roeyers, 2019). Although the findings of altered connectivity in ASD are vast, one finding that is particularly relevant to the current design is that Inflexibility of neural circuitry has been linked to behavioral inflexibility in ASD. For example, it is more difficult to discriminate functional connectivity of specific brain networks (namely, the salience network (SN), default mode network (DMN), and central executive network (CEN)) from each other in ASD relative to typical controls (Uddin et al., 2014). The SN and CEN networks are associated with salient information processing and cognitive control, respectively and nodes include the insula (SN), anterior cingulate (SN), dorsolateral prefrontal cortex (CEN) and posterior parietal cortex (CEN), which have been found to be relevant in performing the current task (see below).

The neural mechanisms of trial-to-trial adjustments and individual differences in learning rate have also been the focus of much recent work. Dynamic fluctuations in learning rate relate to overall arousal levels as measured by pupil diameter (Nassar et al., 2012), as well as activation in a network that includes insula, dorsomedial prefrontal cortex, and parietal cortex, and parts of dorsolateral prefrontal cortex (Behrens et al., 2007; McGuire et al., 2014; Payzan-LeNestour, Dunne, Bossaerts, & O’Doherty, 2013). Functional connectivity over a subgraph that includes many of these regions, and is closely related to both the salience and central executive networks described above, predicts individual differences in learning behavior (Kao et al., 2019). Given the well established connectivity differences in ASD, it is possible both *attention to detail* and adaptive learning in our task are jointly driven by individual differences in functional connectivity, and we hope that our work motivates future explorations of these brain-behavior relationships.

One important question is whether the brain networks that reflect learning rate are actually implementing a learning signal, or doing something more general such as assigning salience to unexpected observations. Two recent studies that clearly dissociate salience from learning using a generative structure like our oddball condition have suggested that the latter may be the case (d’Acremont & Bossaerts, 2016; Nassar, Bruckner, & Frank, 2019a). One recent idea that attempts to rectify differences between the relationship between brain activity and learning rate observed across different statistical environments is that learning rate adjustments are implemented through changes in the active latent state (Nassar, Bruckner, & Frank, 2019a; Nassar, McGuire, Ritz, & Kable, 2019b; Wilson, Takahashi, Schoenbaum, & Niv, 2014). When changes to this latent state are carried forward in time (eg. changepoints) then they could drive increases in learning rate, whereas when they are rapidly replaced (eg. oddballs) then they could drive reductions in learning rate. Within this framework, the fronto-parietal control network could be thought of as providing a signal to load a new state representation, whereas regions including the orbitofrontal cortex seem to reflect the newly loaded state itself (Nassar, McGuire, Ritz, & Kable, 2019b). An important question raised by our work is where individual differences would fall in such a mechanistic process. If the primary determinant of individual differences were in the salience assignments then one might expect divergent individual difference relationships across the changepoint and oddball conditions. However, we observe similar individual difference relationships across the changepoint and oddball conditions, suggesting that *attention to detail* is less related to salience assignments than to the learning itself. This raises important questions about how such individual differences could emerge in the mechanistic model above, and should motivate future neuroimaging studies using individual differences in both task conditions to dissect the neural mechanisms through which *attention to detail* promotes flexible, but unstable, beliefs.

## Conclusions

Our results identify a link between *attention to detail*, a trait elevated in autism, and learning policies that favor flexibility over stability. Individuals high on *attention to detail* pay a price in terms of stability, with beliefs that tend to incorporate fewer observations than they would otherwise. These results were specific to *attention to detail* and unrelated to IQ or other autism linked traits. Overall, our findings demonstrate a core negative consequence of *attention to detail*, namely that by focusing attention on the newest observation, it limits the ability to integrate relevant information across a broader temporal context.

## Open Practices Statement

All data and analysis code associated with this paper will be made available upon acceptance of the publication on the authors website (https://sites.brown.edu/mattlab/resources/). None of the experiments reported here were preregistered.

## Acknowledgments

We thank Ben Heasly for programming the original version of our task. We also thank Antoinette Sabatino DiCriscio and Kayleigh Adamson for assistance with participant testing. This work was supported by a Simons Foundation SFARI Explorer Award (#350225) to V.T. and NIH F32-MH102009-01A1 and K99AG054732 to M.R.N.

## Supplementary Figures

**Figure S1:**
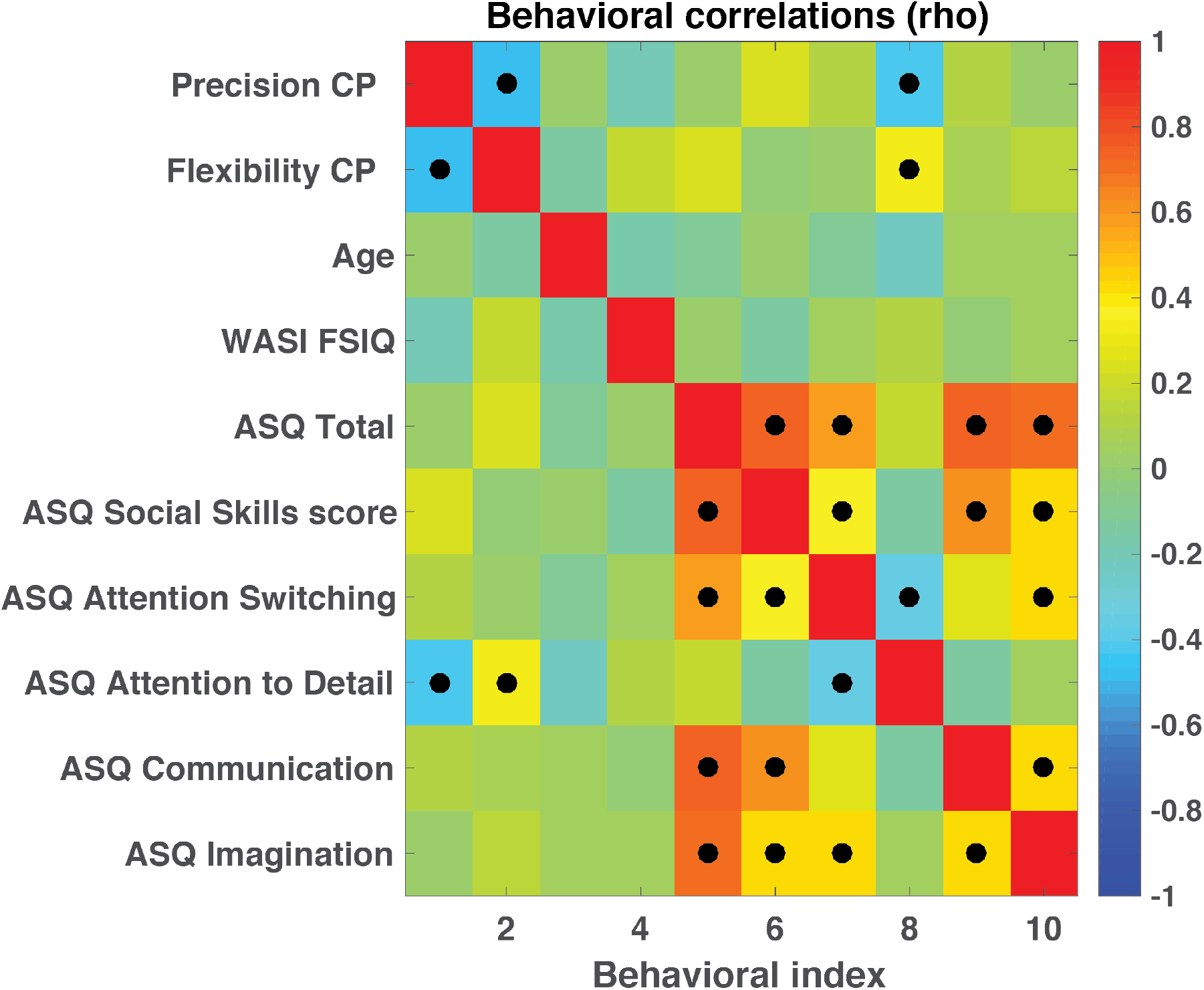
Attention to detail uniquely explains variation in precision and flexibility of beliefs. Color indicates rank-order correlation between each pair of measures and black dots indicate correlations exceeding an uncorrected significance threshold (p < 0.05).

**Figure S2:**
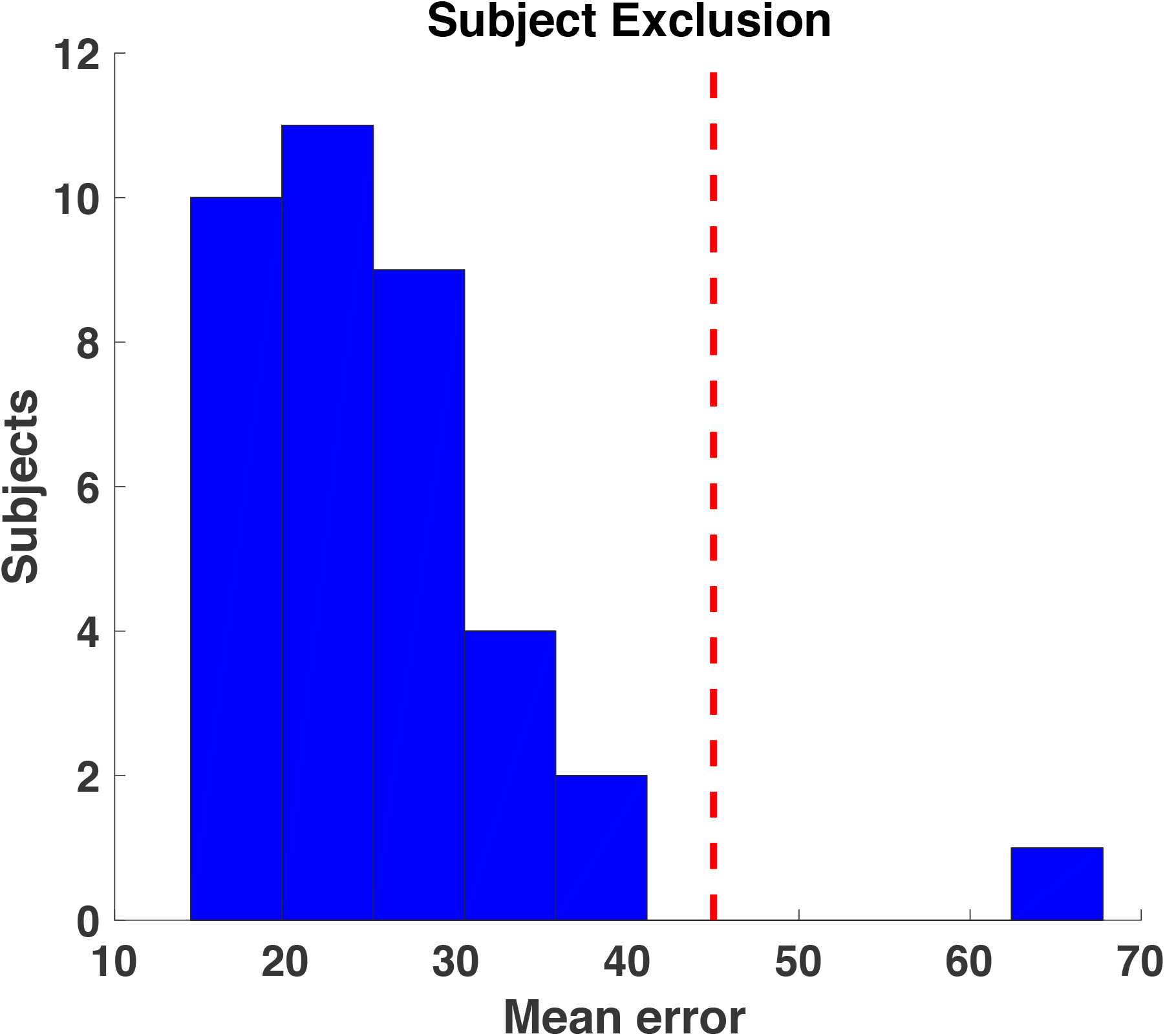
One subject was excluded from the cohort of heterogenous children based on abnormally large mean errors.

**Figure S3:**
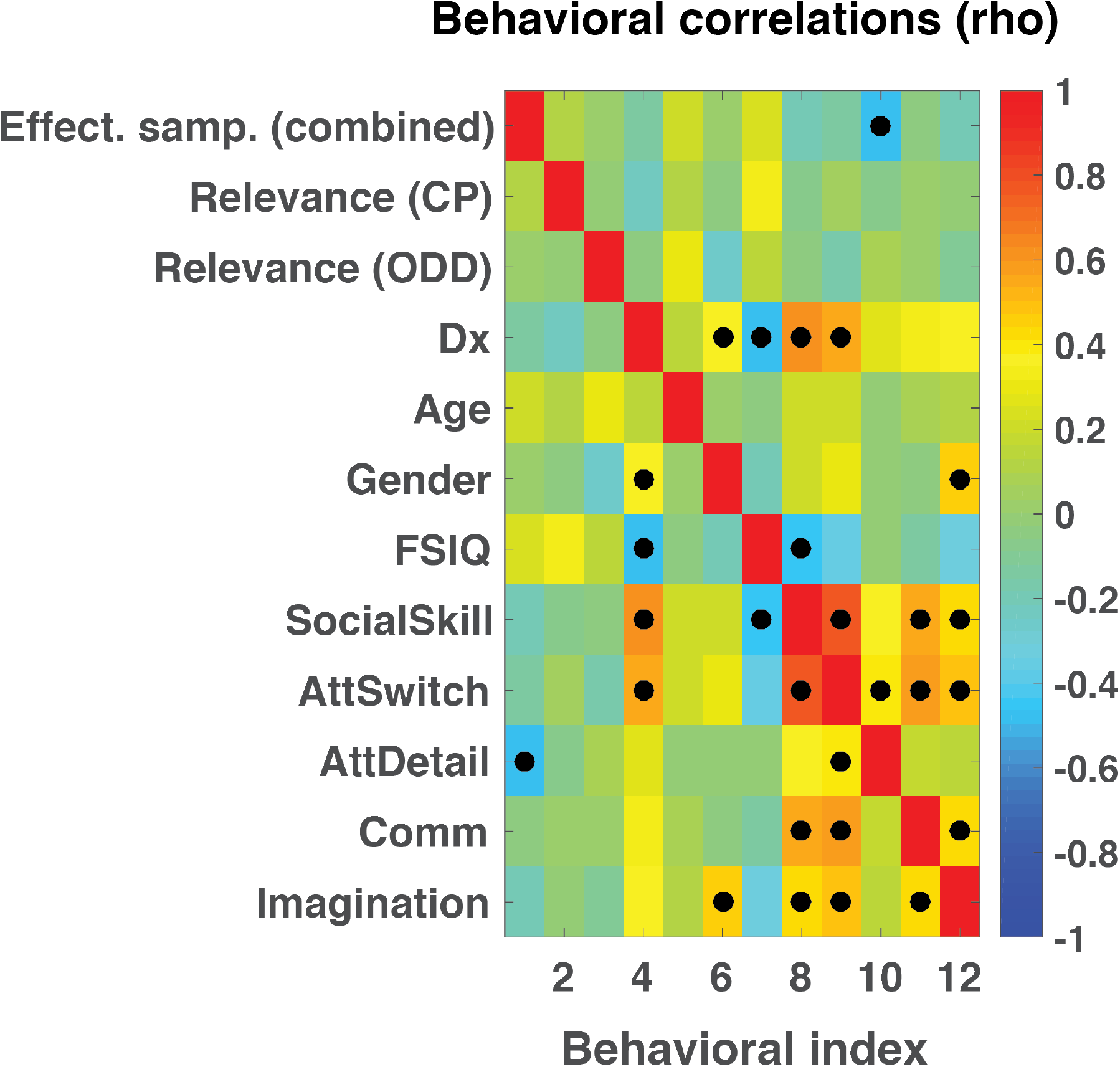
Precision (Effective samples) combined across conditions was negatively related to attention to detail measures from the AQ in the developmental cohort, but not to autism diagnosis category (Dx), Age, Gender, IQ (FSIQ), or other measures from the AQ (SocialSkill, AttSwitch, Comm, Imagination). Color indicates rank-order correlation between each pair of measures and black dots indicate correlations exceeding an uncorrected significance threshold (p < 0.05).

